# Membrane-bounded nucleoid discovered in a cultivated bacterium of the candidate phylum ‘Atribacteria’

**DOI:** 10.1101/728279

**Authors:** Taiki Katayama, Masaru K. Nobu, Hiroyuki Kusada, Xian-Ying Meng, Hideyoshi Yoshioka, Yoichi Kamagata, Hideyuki Tamaki

## Abstract

A key feature that differentiates prokaryotes from eukaryotes is the absence of an intracellular membrane surrounding the chromosomal DNA. Here, we report isolation of an anaerobic bacterium that possesses an additional intracytoplasmic membrane surrounding a nucleoid, affiliates with the yet-to-be-cultivated ubiquitous phylum ‘*Ca*. Atribacteria’, and possesses unique genomic features likely associated with organization of complex cellular structure. Exploration of the uncharted microorganism overturned the prevailing dogma of prokaryotic cell structure.

## Introduction

Cultivation of uncultured microorganisms is a critical step in uncovering their phenotypic features, such as cell structure and metabolic function. However, most lineages of the domains *Bacteria* and *Archaea* remain uncharacterized^1^ due to difficulties in cultivation^2,3^. While omics-based cultivation-independent characterization can circumvent cultivation and provide insight into their metabolism and ecology^4,5^, metabolic reconstruction is generally based on genes characterized in cultured organisms and, thus, prediction of novel phenotypic features of uncultured microorganisms remains challenging^6^.

## Results and Discussion

In this study, we succeeded in isolating a novel anaerobic bacterium (pointed rod-shape and non-spore-forming), designated strain RT761, that belongs to the clade OP9^7^ of ‘*Ca*. Atribacteria’^5^ and possesses double-layered intracytoplasmic membrane (ICM) (Fig. 1), after 3 years of enrichment from saline formation water and sediments derived from deep aquifers in natural-gas deposits in Japan. The ICM clearly compartmentalizes the cytoplasm into a nucleoid-present space (referred to as ICM-bound space, IBS) and - absent space (referred to as cytoplasmic membrane-bound space, CBS) (Fig. 1c, 1d), envelopes the nucleoid during the entire course of cell division and is split into the daughter cell as division complete (Supplementary Fig. S1).

**Fig. 1.**
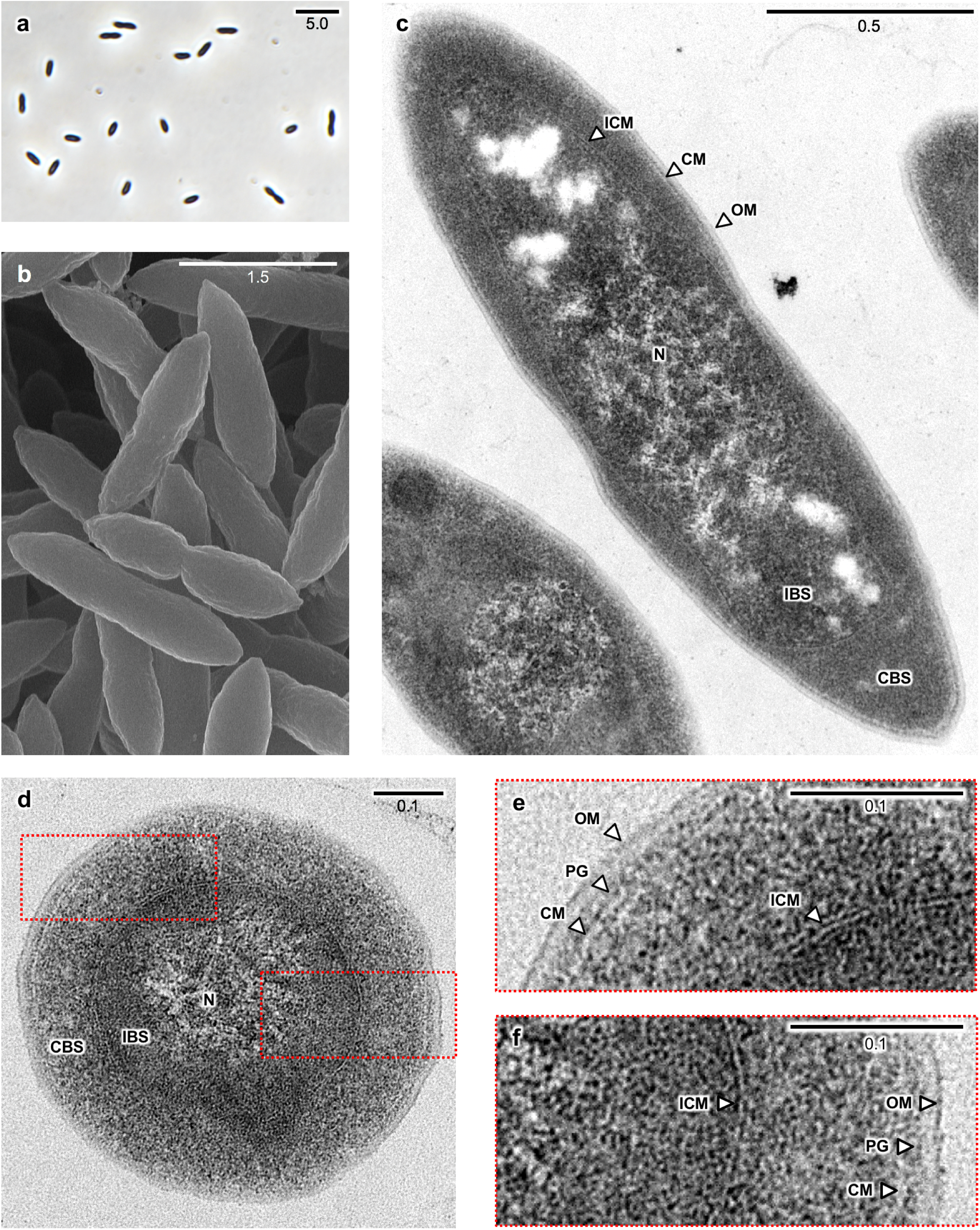
Morphology and membrane structure of RT761 cells showing the presence of intracytoplasmic membranes surrounding the nucleoid. Phase-contrast (**a**) and scanning electron (**b**) microscopy showed a pointed rod shape of RT761 cells. Transmission electron microscopy (**c-f**) showed a gram-negative cell structure consisting of an outer membrane (OM), thin peptidoglycan (PG)-like layer, cytoplasmic membrane (CM) and an additional intracytoplasmic membrane (ICM). Abbreviation: CBS, CM-bound space; IBS, ICM-bound space; N, nucleoid. (Scale bars: μm.)

Using confocal laser scanning microscopy, we observed a distinct space in RT761 cells between the outer rim of the cells and DNA/RNA along with a lipid membrane structure that appears to define this boundary, which most likely corresponds to the ICM (Fig. 2). Ribosomes were also observed to be bound by this ICM using fluorescence *in situ* hybridization with an rRNA-targeted probe (Supplementary Fig. S2 and Supplementary Discussion 1). These results indicate that DNA replication, transcription, and translation take place mainly in the IBS. Although a member of the *Planctomycetes* phylum, *Gemmata obscuriglobus*, was also thought to form intracytoplasmic membrane surrounding nucleoid in a cell^8^, a recent study indicated that what appeared to be an ICM of planctomycetal cells was invagination of the CM^9^. Thus, RT761 is the first bacterium that can form an unusual subcellular lipid bilayer-bound structure that contains genetic materials and participates in core genetic processes. Remarkably, membrane potential (ΔΨ) across both the CM and ICM was detected using a ΔΨ-sensitive dye (3,3’-dihexyloxacarbocyanine iodide [DiOC_6_]^10^) (Supplementary Fig. S3), suggesting energy metabolism/consumption in both CM/CBS and ICM/IBS. Moreover, examples of subcellular lipid membrane-bound structures are scarce across the two prokaryotic domains (*e.g.*, magnetosomes, anammoxosomes, photosynthetic membranes like chromatophores and thylakoids)^11^, further highlighting the uniqueness of this finding.

**Fig. 2.**
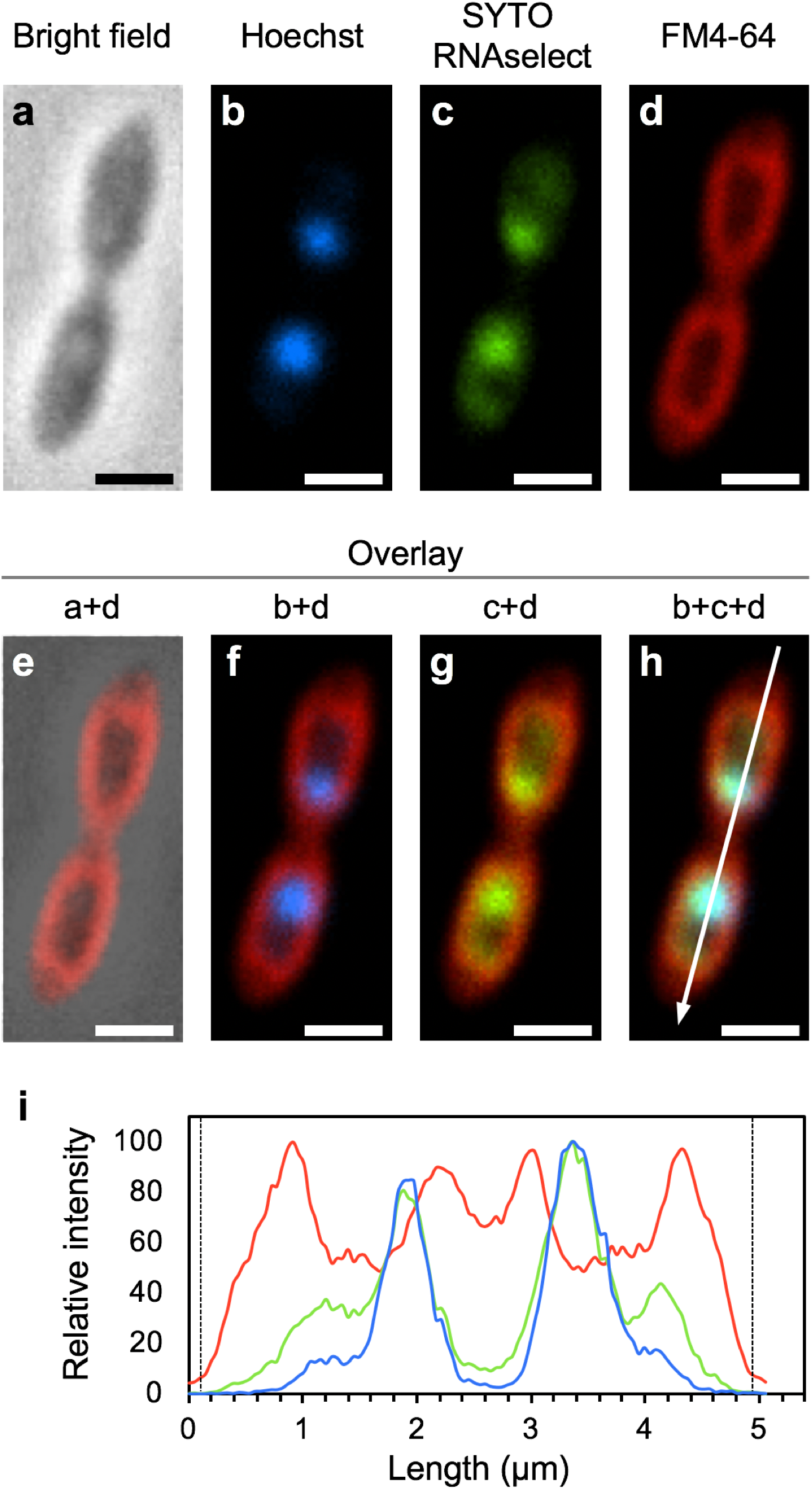
Confocal-laser microscopy showing the presence of intracytoplasmic membranes (ICM) and localization of DNA and RNA within the RT761 ICM. DNA, RNA and membrane lipids were stained by Hoechst (blue), SYTO RNAselect (green) and FM4-64 (red) respectively. (**a**), Phase contrast image. (**b-d**) Confocal-laser images. (**e-h**) Image overlays. (**i)** Line profiles of fluorescence intensity plotted longitudinally along white arrow in (h). Broken lines indicate the edges of cell observed in bright field (a). (Scale bars: 1 μm.)

In both dividing and non-dividing cells, RNA not only localized with chromosomal DNA, but also co-localized at the polar ends of the IBS (Fig. 2 and Supplementary Fig. S4). This coincides with the section of ICM that separates the IBS and the largest region of CBS. While subcellular localization of RNA of specific genes has been observed in bacterial species^12^, the localization of bulk RNA has not. Since mRNAs can localize to specific regions in bacterial cells where their protein products function^13^, we speculate that the ICM section that separates IBS and CBS may play an important role in regulating RT761’s physiology.

The importance of localization and membranes in RT761 was further supported by genomic and transcriptomic analyses. Alignment of all RT761 protein-coding genes with a reference sequence database revealed that 34 genes in the RT761 genome contained unique N-terminal extensions (NTE; 10-73 amino acids in length) compared to the top 250 hits in the NCBI RefSeq database and some of them were conserved among ‘*Ca.* Atribacteria’ OP9 genomes (Supplementary Table S1 and Table S2). The genes with NTE included those involved in critical cellular processes: cell division (FtsZ), Lipid A biosynthesis (UDP-3-O-acyl-*N*-acetyglucosamine deacetylase – LpxC), DNA replication, DNA repair, transcriptional regulation, tRNA processing, transmembrane signaling, and H_2_ generation (FeFe hydrogenase subunit alpha – HydA). Notably, several facilitate central functions in their respective processes: FtsZ recruits other cell division proteins to the fission site^14^, LpxC performs the committing step in Lipid A biosynthesis^15^, and HydA catalyzes the reduction of protons to H_2_ in the hydrogenase complex^16^. NTEs in prokaryotes have so far been only found in enzymes that localize to the lumen of subcellular compartments called bacterial microcompartment (BMC) and are necessary for the encapsulation of enzymes with NTE into BMC shells^17,18^. Among 34 genes with NTE in RT761, only one enzyme (deoxyribose-phosphate aldolase) is expected to localize to the BMC^4^. Given the observation of DNA and RNA localization, we speculate that some of NTEs are signal sequences for subcellular localization, which is a necessary feature for RT761 to regulate cell function within its complex cellular structure. Interestingly, RT761 and other ‘*Ca*. Atribacteria’ possessed two FtsZ genes, one with an NTE and the other without (Supplementary Fig. S5). RT761 expressed both FtsZ’s during exponential growth. FtsZ is known to localize to the membrane through interaction with cell division protein FtsA that associates with the membrane through a C-terminal amphipathic helix^19^. While the NTE-lacking FtsZ gene is adjacent to FtsA, the FtsZ with an NTE lacks a corresponding FtsA. The *Ca*. Atribacteria FtsZ NTE were predicted to form amphipathic helices that can bind to the membrane (Supplementary Fig. S5), suggesting that the two FtsZ have a different localization mechanism and non-redundant roles in RT761. Although 9 out of 6,751 cultured bacterial type strains possess both a typical and NTE-possessing FtsZ, putative amphipathic helices were not found in any of these sequences. Such unique features may be essential for binary fission through triple lipid membranes in RT761 (Supplementary Fig. S1).

Further analysis reveals unique genomic features of RT761 related to membrane-mediated physiology. Based on transcriptomic analysis of RT761 under exponential growth phase, membrane-associated proteins comprised 5 out of 10 of the mostly highly expressed genes (Supplementary Table S3). These include a putative transmembrane protein, lipoprotein, periplasmic substrate-binding protein, and two fasciclin domain-containing transmembrane proteins, all of which have unknown functions. These findings point towards importance of membrane-centric metabolism in RT761 physiology. The RT761 genome also has a high proportion of proteins with transmembrane helices (29.6% of all proteins) greater than 99.7% of all gram-negative type strains with sequenced genomes available (Supplementary Fig. S6). We also found that RT761 may have unique signal peptide sequences for Sec-secreted proteins through comparison of results from different algorithms. While SignalP-4.1^20^ estimated that RT761 has a low proportion of Sec-secreted proteins (3.4% of all proteins) less than 96.7% of all gram-negative type strains, SignalP-5.0^21^ predicted 2.67 times more (9.0% of all proteins) (Supplementary Fig. S6). Evaluation of all gram-negative type strain genomes revealed that most cultured phyla (26 out of 29) have consistent predictions between SignalP-4.1 and SignalP-5.0 (1.1 ± 0.2 [S.D.] times more on average) (Supplementary Discussion 2); remarkably, we only observed RT761-like signatures (high genomic proportion of proteins with transmembrane helices and underestimation of Sec-secreted proteins by SignalP-4.1) in three other cultured phyla with unique cell structures (Supplementary Fig. S6): *Thermotogae* members (outer toga^22^), *Dictyoglomi* (multi-cell-spanning outer envelope^23^), and *Caldiserica* (electron-lucent outer envelope^24^). In total, comparison of genomic transmembrane and extracellular protein abundance signatures may serve as a new approach for identification of bacterial lineages with novel cell membrane structure and is, thus, quite distinct from currently available genotype-based cell morphology prediction approaches (*e.g.*, RodZ for rod-shape and lipid A synthesis genes for gram-negative structure).

In addition to the unique cell structure and genomic feature, we found that strain RT761 is capable of syntrophic interaction. RT761 fermented glucose, producing H_2_, acetate, CO_2_ and ethanol (trace levels) as end products and could not utilize exogenous electron acceptors for anaerobic respiration (*i.e*., nitrate, ferric iron, and sulfate). Although RT761 growth was inhibited by accumulation of hydrogen during cultivation with glucose, addition of a hydrogen-consuming methanogenic archaeon significantly increased the growth rate and maximum cell density of RT761 (Supplementary Fig. S7). RT761 can theoretically shift to ethanol fermentation as an alternative electron disposal route but only generates a small amount (Supplementary Fig. S7), indicating that RT761 primarily relies on hydrogen formation to maintain cellular redox balance. Thus, in contrast to most hydrogen-producing fermentative bacteria, RT761 highly depends on syntrophic association with hydrogen-scavenging methanogen for ideal growth. Such dependence of sugar degradation on a syntrophic partner is thought to be important in anoxic ecosystems^25,26^. We speculate that RT761 may also avoid ethanol production as continuous exposure to acetaldehyde generated through ethanol fermentation could cumulatively damage chromosomal DNA, especially due to the slow growth rate (doubling time of 5.1 days). In addition, we observe the expression of the gene cluster encoding homologues of BMC previously proposed to sequester and condense aldehydes^4^. Similar metabolisms of BMC-mediated aldehyde conversion to sugars and syntrophy that are theoretically possible or thermodynamically required for association with methanogens have been predicted in cultivation-independent analyses of ‘*Ca*. Atribacteria’^4,5,27,28^. The observed syntrophic lifestyle of RT761 justifies the prevalent detection of environmental clones of ‘*Ca*. Atribacteria’ across Earth’s anoxic ecosystems favoring fermentation and syntrophy^4^.

Phylogenetic analysis based on 16S rRNA gene and conserved protein-coding markers revealed that strain RT761 was assigned to the clade OP9 of ‘*Ca*. Atribacteria’ (Supplementary Fig. S8 and S9), making this strain the first culturable representative of this candidate phylum since its 16s rRNA-based discovery in sediments from the hot spring in Yellowstone National Park and designation as OP9 in 1998^7^. Based on phenotypic, genotypic and phylogenetic characteristics, we propose strain RT761 as a new species, ‘*Ca*. Atrimonas tricorium’ (A.tri.mo’nas. L. adj. *ater -tra -trum*, black; L. fem. n. *monas*, a unit; N.L. fem. n. *Atrimonas*, a bacterium isolated from the dark, deep sedimentary environment) (tri.co’ri.um. L. pref. *tri*, three; L. neut. n. *corium*, layer or coating; N.L. neut. n. *tricorium*, triple membrane).

We discovered a bacterium belonging to a hitherto uncultivated ubiquitous phylum whose cell structure, organization and regulation are much more complex than a typical prokaryote, providing a new perspective on prokaryotic cell biology. Further characterization of *‘Ca*. A. tricorium’ may help us uncover the evolution of the prokaryotic ability to form intracellular membranes surrounding chromosomal DNA and its relationship to development of the eukaryotic nucleus.

## Acknowledgments

We acknowledge the Kanto Natural Gas Development Co., Ltd. for collecting environmental samples at their facilities. We thank Naoki Morita for quantification of fermentation products; Chiwaka Miyako for assistance in molecular analyses; Fumie Nozawa for assistance in cultivation experiments. This work was supported by JSPS KAKENHI Grant Numbers JP17K15183, JP18H05295 and JP18H02426.

## Author Contributions

T.K., M.K.N., Y.K., and H.T. designed the study and wrote the manuscript. T.K. and H.Y. performed enrichment cultures, and T.K. isolated the bacterium. M.K.N. performed bioinformatic analyses. T.K., H.K. and X.Y.M. performed microscopic analyses. All authors reviewed the results and approved the manuscript.

## Author Information

The draft genome sequences and annotation data of strain RT761 are available in NBCI BioProject under accession number PRJNA528842. The authors declare no competing financial interests. Correspondence and requests for materials should be addressed to H.T. and Y.K.

