## Supplementary Information for "Membrane-bounded nucleoid discovered in a cultivated bacterium of the candidate phylum ‘Atribacteria’"

**This file includes:**

**Methods**

**Supplementary Figs. S1 to S10**

**Supplementary Discussions 1 and 2**

**References**

### Methods

#### Sample collection

The sediment and formation water samples were collected from a settling pond that was placed downstream of a commercial gas and water producing well to remove suspended sand particles from the formation water in Mobara, Chiba prefecture, Japan. The samples came from the gas-bearing aquifers in the screened depth range of 490-900 m that consist of repeating sequences of turbidite (alternating beds of sandstone and mudstone) in the Otadai and Kiwada formations. These sediments were deposited in deep marine environments during the Plio-Pleistocene periods[29,30]. The water temperature was 24.4°C, the pH was 7.7, and the redox potential was -213 mV. The $Cl^-$ concentration was 17,000 mg $l^{-1}$, and the sulfate concentration was <5 mg $l^{-1}$ (detection threshold). The natural gases produced in this area composed mainly of methane (99%), and the origin of methane was suggested to be of biogenic based on stable isotopic analysis[31].

The samples were collected in sterilized glass bottles with butyl rubber stoppers and screw caps. The bottle was purged with $N_2$ gas prior to sample collection and was filled with the water to maintain the samples under anaerobic conditions.

#### Enrichment culture and isolation

Sediment samples were mixed at a 1:2 volume ratio with a formation water to make slurry samples in an anaerobic chamber. The slurry samples were dispensed as 20 ml-aliquots into 75-ml serum vials and were then sealed using butyl rubber stoppers and aluminum crimps in an anaerobic chamber. The slurries were incubated without the addition of any nutrients under an atmosphere of $N_2/CO_2$ (80: 20) at a temperature higher than the water temperatures of original environments (45 °C rather than 25 °C). After 90 days, 2 ml of the methane-producing culture of slurry sample was inoculated into a saline mineral medium[32] containing 1 g $l^{-1}$ glucose, 1 g $l^{-1}$ Bacto peptone (BD), 0.1 g $l^{-1}$ yeast extracts

(BD), 5 mM coenzyme M (2-mercaptoethane sulfonic acid) and 0.1 mM titanium (III) citrate (used as a reducing agent). Cultivation was performed in 75-ml serum vials containing 20 ml of medium under an atmosphere of $N_2/CO_2$ (80: 20). Enrichment cultures were grown, followed by successive transfer six times at intervals of approximately 80 days. Individual cells were isolated in a pure culture using the deep agar slant method combined with dilution-to-extinction method[32] with a saline mineral medium supplemented with 1 g $l^{-1}$ glucose, 0.1 g $l^{-1}$ yeast extracts, 0.1 mM titanium (III) citrate and 8 g $l^{-1}$ agar. After 40 days of incubation, a single colony was picked and transferred to fresh liquid medium. This procedure was repeated three times. Purity of the culture was verified by microscopy and further confirmed by no contaminant sequences in DNA sequencing data of genomic DNA. The pure culture of strain RT761 was incubated at 45°C in saline mineral medium amended with 16 mM glucose, 0.2 g $l^{-1}$ yeast extracts and 0.5 g $l^{-1}$ cysteine hydrochloride. In co-culture with methanogenic archaeon, *Methanothermobacter thermoautotrophicus* strain Delta H, 5 mM coenzyme M and 0.2 g $l^{-1}$ $Na_2S\cdot 9H_2O$ were also added.

**Genomic and transcriptomic analyses**

Genomic DNA of strain RT761 was extracted using the Blood and Cell Culture DNA Maxi Kit (QIAGEN, Venlo, Netherlands) according to the manufacturer's instructions. DNA was sequenced using a PacBio RS II system (PacBio, Menlo Park, CA) with one single-molecule real-time (SMRT) cells at Hokkaido System Science (Sapporo, Japan). Sequence assembly was carried out using the GS De Novo assembler Newbler (version 2.3). Gene identification and annotations were performed using annotated by Prokka v1.13[33]. To search for unique N-terminal extensions, all RT761 protein-coding genes were aligned with the Genbank RefSeq database[34] using BLASTP[35] and compared with all top 250 BLASTP hits filtered with >30% similarity and >70% coverage. Secondary structure of amino acid sequences of N-terminal extension associated with FtsZ was predicted using JPred4[36]. Transmembrane proteins and signal peptides were predicted using TMHMM v2.0[37] (default options) and SignalP (v4.1[20] and v5.0[21] using the gram-negative option and default options for the remaining settings) correspondingly, and the percentages of these proteins out of the total number of ORFs were compared to those in all gram-negative type strain draft genomes available on the Joint Genome Institute Integrated Microbial Genomes and Microbiomes database[38] and those in draft genomes of uncultured phyla that were confirmed to encode lipid A synthesis genes (lpxB, lpxC, or lpxD) in at least one draft genome. The sequences were aligned against the SILVA v132 alignment using SILVA[39] with default settings. The phylogenetic tree was constructed using RAxML-NG[40] using the generalized time reversible (GTR) model, 4 gamma categories, and 100 bootstrap iterations. Prediction/selection of conserved genes and tree construction was performed through PhyloPhlAn[41] using default settings.

RNA was extracted from late exponential growth of both pure culture and co-culture with a methanogen *M. thermoautotrophicus* str. Delta H using the ISOSPIN Plant RNA kit (NIPPON GENE, Japan) according to the manufacturer's instructions and was sequenced using an Illumina sequencer NovaSeq 600 system (illumina, USA) at Filgen, Inc. (Nagoya, Japan). Total RNA was depleted of ribosomal RNA via Ribo-Zero rRNA removal kit (illumina). The sequenced RNA was trimmed via Trimmomatic v0.33[42] and mapped to the assembled genome through BBmap v37.10 (https://sourceforge.net/projects/bbmap/) to calculate the gene expression levels, which were represented by Reads Per Kilobase of transcript per Million mapped reads.

### Morphological characterization

Cell morphology was observed via phase-contrast and fluorescence microscopy (BX51; Olympus), confocal laser scanning microscopy (LSM800; ZEISS), scanning electron microscopy (SEM) (S-4500; Hitachi) and transmission electron microscopy (TEM) (H-7600; Hitachi). The cells in exponential phase of growth in pure culture condition were used for all microscopic observation.

Cells were washed with phosphate-buffered saline (PBS) before staining. Membranes of RT761 cells were strained with FM4-64 (ThermoFisher Scientific) at a final concentration of 40 μg $ml^{-1}$. DNA was strained with SYBR Green I or Hoechst 33342 (ThermoFisher Scientific) at final concentrations of 1-2 μg $ml^{-1}$. RNA was strained with SYTO RNAselect (ThermoFisher Scientific) at a final concentration of 10 μM. To visualize membrane potential[10], cell membranes were stained with $DiOC_6(3)$ (ThermoFisher Scientific) at a final concentration of 5 μg $ml^{-1}$. The stained sample was incubated for 1 h at 30°C and observed under phase-contrast or confocal laser scanning microscopes.

For SEM observation, the cells were fixed with 2% glutaraldehyde in 0.1 M sodium phosphate buffer (pH7.2) at 4°C for 2 h, postfixed with 1% osmium tetroxide at room temperature for 1h, dehydrated through a graded ethanol series followed by 3-methylbutyl acetate for 20 min, dried with a critical point dryer (JCPD-5; JEOL), and finally coated with gold.

For TEM observation, the cells were fixed with 2.5% glutaraldehyde in 0.1 M sodium cacodylate buffer (pH7.4) at 4°C for 3 h and then postfixed with 1% osmium tetroxide at 4°C for 90 min. The fixed cells were suspended in 1% aqueous uranyl acetate at room temperature for 1 h. The suspended cells were embedded in 1.5% agarose and dehydrated through a graded ethanol series. The dehydrated blocks were embedded in Epon812 resin. Ultrathin sections were cut with an ultramicrotome (Leica EM UC7), mounted on copper grids, and stained with uranyl acetate and lead citrate.

For fluorescence *in situ* hybridization, the cells were fixed in 1% paraformaldehyde at 4°C for overnight and stored in 99% ethanol-PBS (1:1) at -20°C. The fixed cells were incubated in a moisture chamber with a hybridization buffer (0.9 M NaCl, 0.01% sodium dodecyl sulfate, 20 mM Tris-HCl, pH 7.2 containing fluorescently labeled probes (0.5 pmol $\mu l^{-1}$). After incubation at 46°C for 2.5 h, the buffer was replaced with washing solution (0.9 M NaCl, 0.01% sodium dodecyl sulfate, 20 mM Tris-HCl, pH 7.2). The sample was incubated at 48°C for 30 min and observed under a fluorescence phase-contrast microscope. An oligonucleotide probe targeting the 16S rRNA gene was Cy-3-labeled EUB338 probe (5'- GCTGCCTCCCGTAGGAGT-3').

**Quantification of metabolites**

Hydrogen and methane in gas phase of cultures were measured with a gas chromatography equipped with a thermal conductivity detector (GC-8A; Shimadzu). Glucose, acetate and ethanol in liquid phase of cultures were measured with a high-performance liquid chromatography (HPLC) (LC20; Shimadzu) with Shim-pack SPR-H column (Shimadzu) or HPLC (LC-2000Plus, Jasco) with Aminex HPX-87H column (BIO-RAD).

**Quantitative PCR**

SYBR green-based real-time PCR was run on a CFX Connect real-time PCR detection system (Bio-Rad Laboratories Inc., Hercules, CA) using the PowerUp SYBR green master mix (Applied Biosystems, California, USA) to quantify the population of RT761 cells. The forward and reverse primers, rt1F (5’-GCTAATACCCCATATGCTCCCTG-3’) and rt1R (5’-ACCTCGCCAACCAGCTGATGGGG-3’), were designed from the 16S rRNA gene sequences of strain RT761. The length of amplified products was 62 bp. Total DNA was extracted from pure- or co-cultures using a ISOSPIN Fecal DNA (NIPPON GENE). Standard curves for quantification were determined based on 10-fold serial dilutions of the target PCR products of strain RT761 at known concentrations. All reactions, including the non-template control, were performed in triplicate. The presence of a single PCR product without any nonspecific amplicons was confirmed via agarose gel and melting curve analyses. The PCR product was sequenced by Sanger sequencing to confirm the amplification of 16S rRNA gene from strain RT761. All qPCR runs showed no PCR amplifications from non-template control and culture samples without adding RT761 cells, and had efficiency levels of approximately 95%, with an $R^2$ of >0.99. Cell growth rate was estimated using 16S rRNA gene copy number as a proxy for cell population.

**Supplementary Figures**

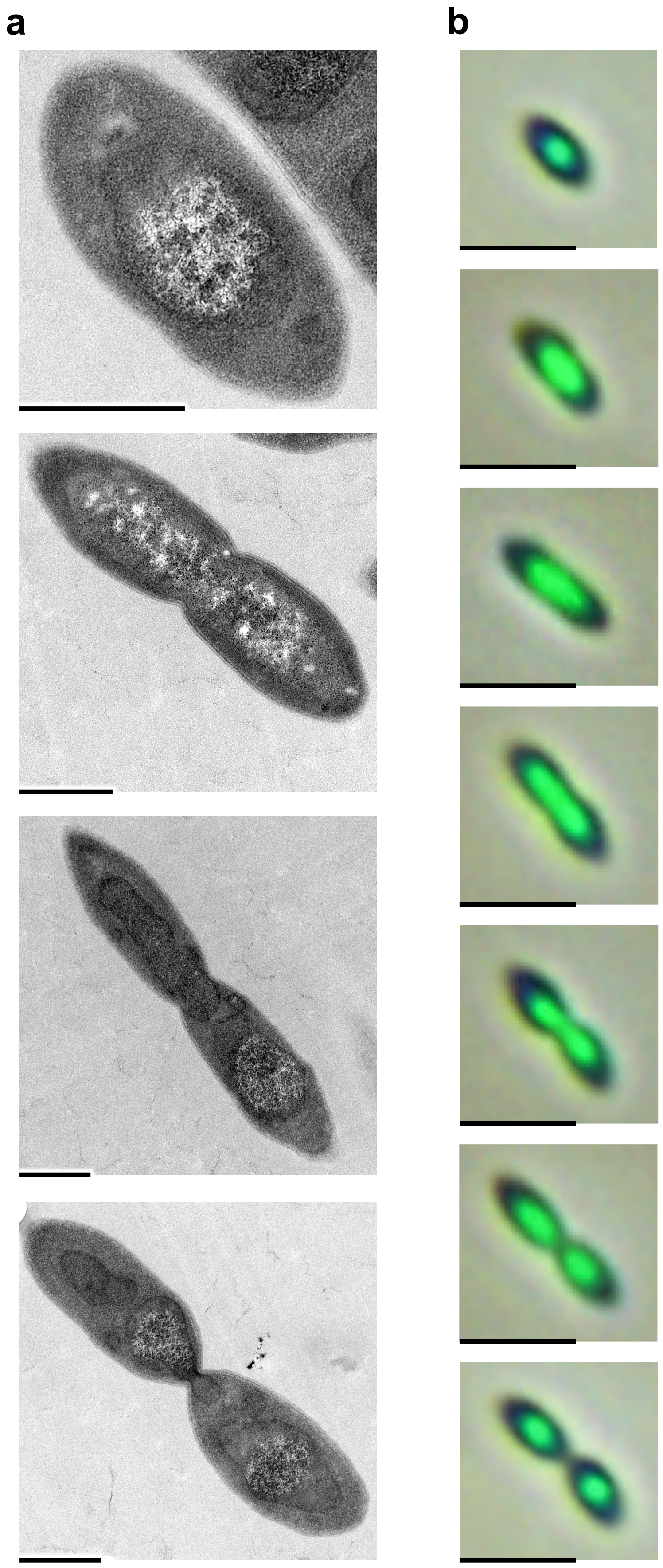


**Fig. S1.** The appearance of intracytoplasmic membrane (ICM) and nucleoid during cell division. (**a**) Transmission electron micrographs showing the occurrence of nucleoid inside the ICM during cell division. (**b**) Light and fluorescence micrographs using a DNA staining dye (SYBR Green I) showing the localized chromosomal DNA during cell division.
Scale bars: 0.5 μm (a) and 2 μm (b).

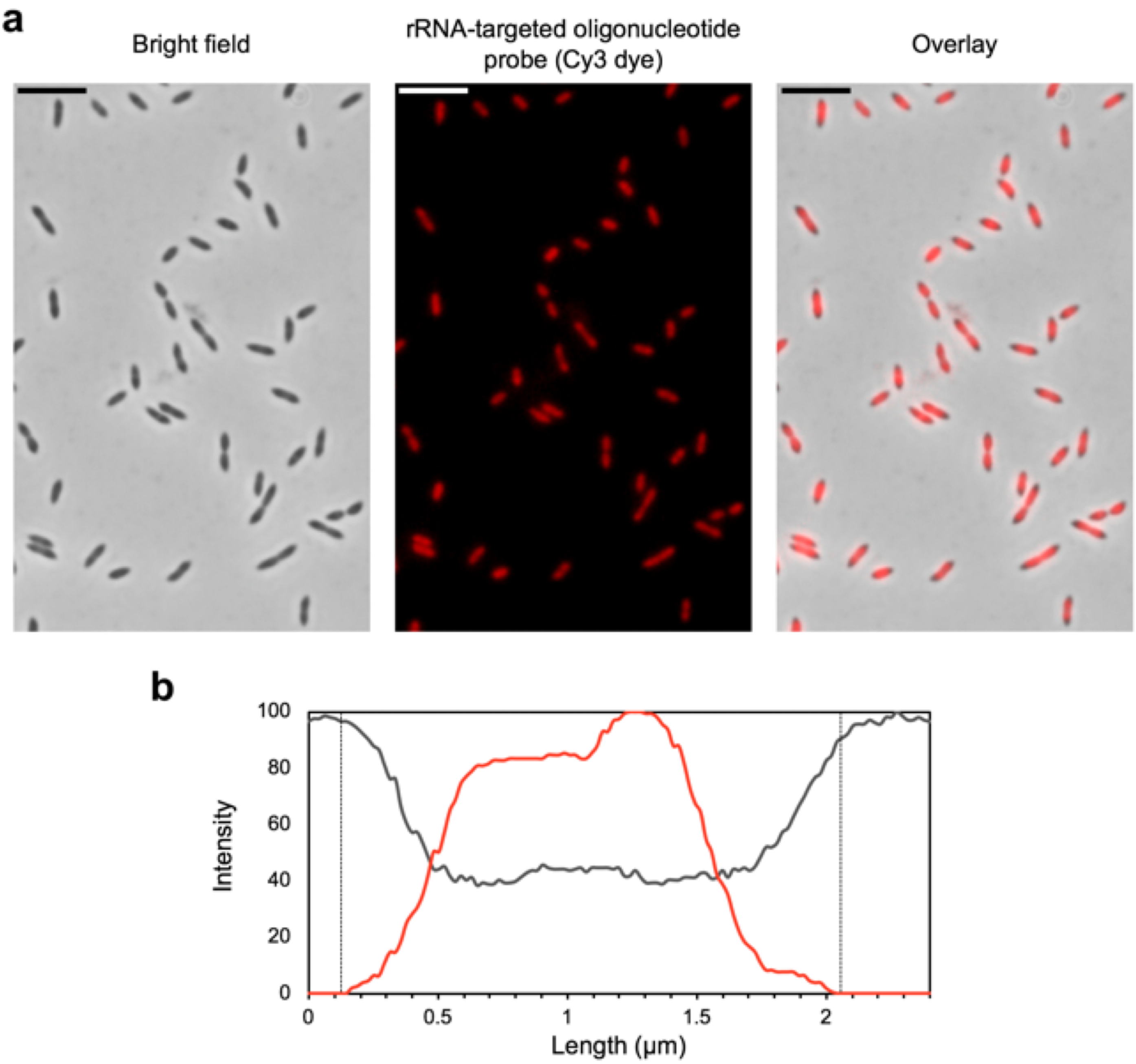


**Fig. S2.** 16S ribosomal RNA (rRNA) staining of formamide-fixed cells using fluorescence *in situ* hybridization showing the localization of ribosomes in RT761 cells. (**a**) Phase contrast micrographs. (**b**) Line profiles of signal intensity of cell (black) and rRNA (red). Broken lines indicate the edges of cell observed in bright field. (Scale bars: 5 µm.)

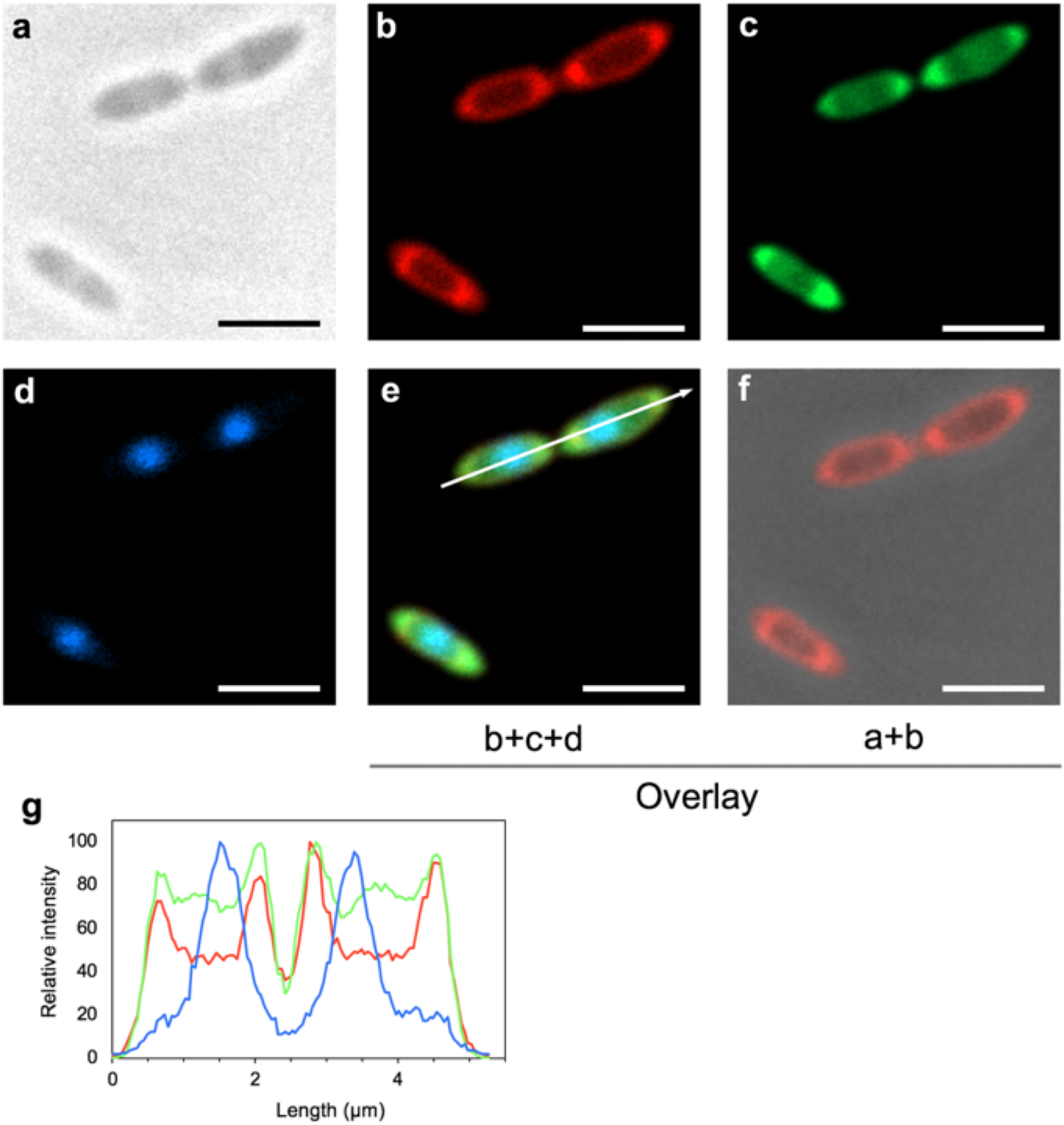


**Fig. S3.** Confocal-laser microscopy showing the membrane potential in both cytoplasmic and intracytoplasmic membranes of RT761 cells. Membrane lipids, energized membrane and DNA were stained by FM4-64 (red), $DiOC_6$ (green) and Hoechst (blue) respectively. (**a**) Phase contrast image. (**b-d**) Confocal-laser images. (**e, f**) Image overlays. (**g**) Line profiles of fluorescence intensity along the white arrow show consistency between FM4-64 (cytoplasmic and intracytoplasmic membranes) and $DiOC_6$ (energized membrane). Lipid membranes in the outer rim of the cells and inside the cells are shown in the overlay picture (**f**). (Scale bars: 2 μm.)

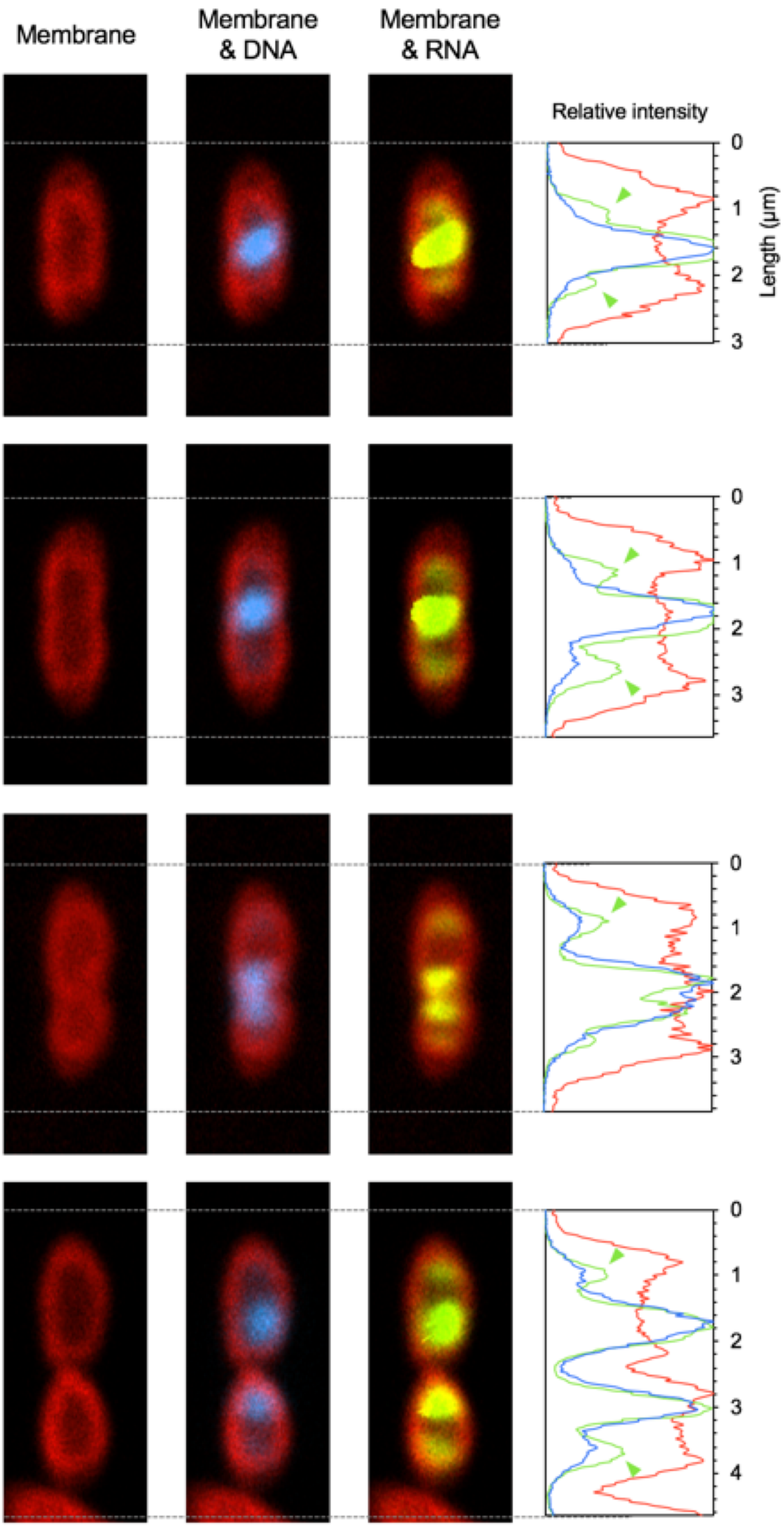


**Fig. S4.** Confocal-laser microscopy showing the localization of RNA during cell division. Membrane lipids, DNA and RNA were strained by and FM4-64 (red), Hoechst (blue) and SYTO RNAselect (green) respectively. Signal peaks for RNA that do not overlap with those of DNA are indicated with green arrowheads.

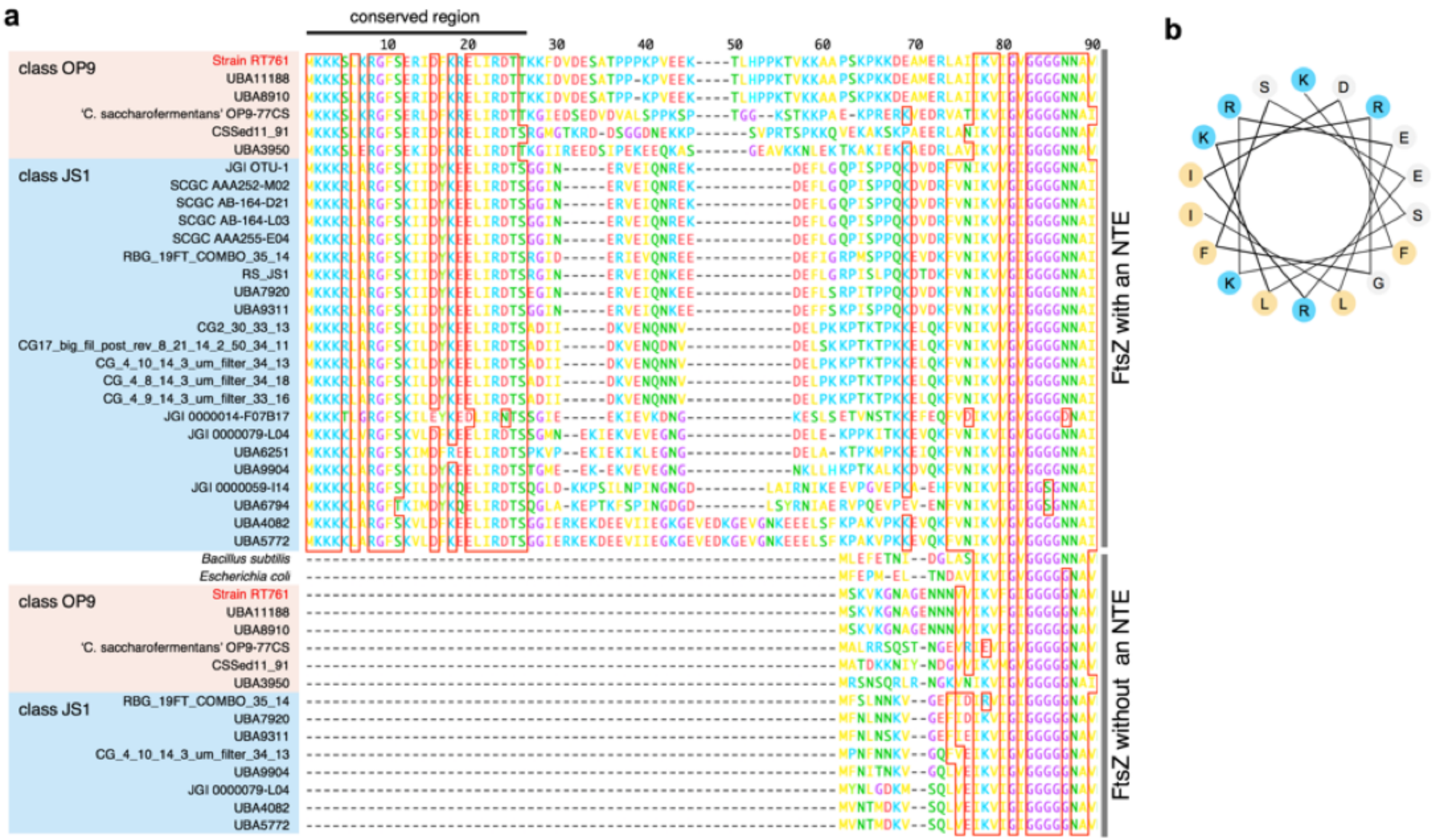


**Fig. S5.** The N-terminal extensions (NTEs) of FtsZ in '*Ca*. Atribacteria' containing a conserved region that has the capacity to form an amphipathic helix. (**a**) Comparison of the *N*-terminal amino acid sequences of FtsZ with and without an NTE from '*Ca*. Atribacteria' genomes. In RT761, two FtsZ proteins shared 86% amino acid sequence similarity. (**b**) Helical wheel diagram of the conserved region from FtsZ with an NTE of RT761. Large hydrophobic and basic residues are highlighted in yellow and blue respectively.

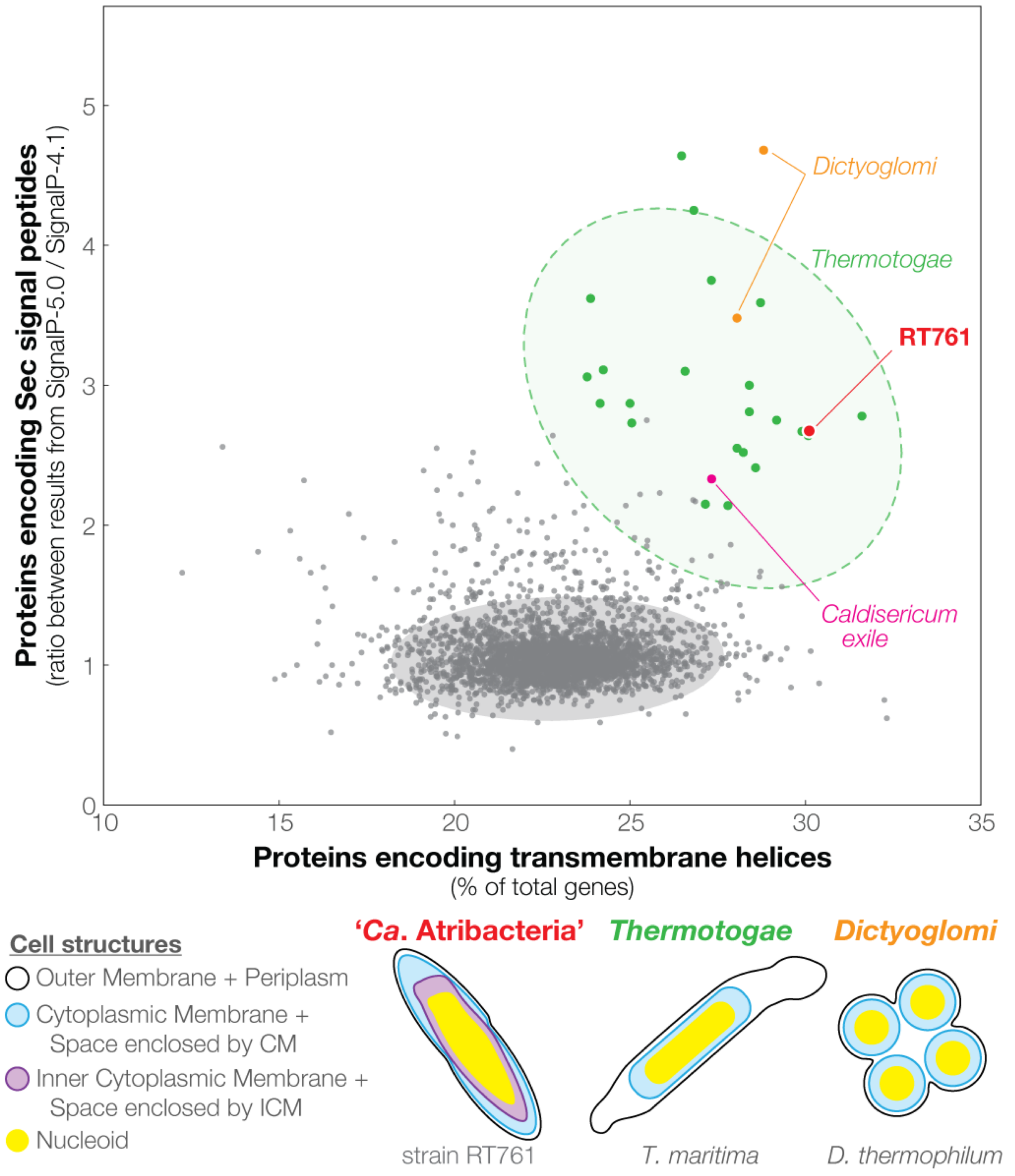


**Fig. S6.** Unique genomic compositions of membrane-related features observed for phyla with unique cell structures. The horizontal axis shows the genomic proportion proteins encoding transmembrane helices. The vertical axis shows the ratio of proportions of proteins encoding Sec signal peptides estimated by SignalP-5.0 and SignalP-4.1. RT761 (red), type strains from *Thermotogae* (green), *Dictyoglomi* (orange), *Caldiserica* (pink), and other gram-negative type strains (gray) are plotted (3,502 genomes downloadable from the Joint Genome Institute Integrated Microbial Genomes and Microbiomes database). For *Thermotogae* and other gram-negative type strains, 95% (green) and 99.9% (gray) confidence ellipses are shown respectively. (Bottom) Cell structures of select species are shown for '*Ca.* Atribacteria', *Thermotogae*, and *Dictyoglomi*. The illustrations indicate the outer membrane (black), cytoplasmic membrane (blue), inner cytoplasmic membrane (purple), and nucleoid (yellow).

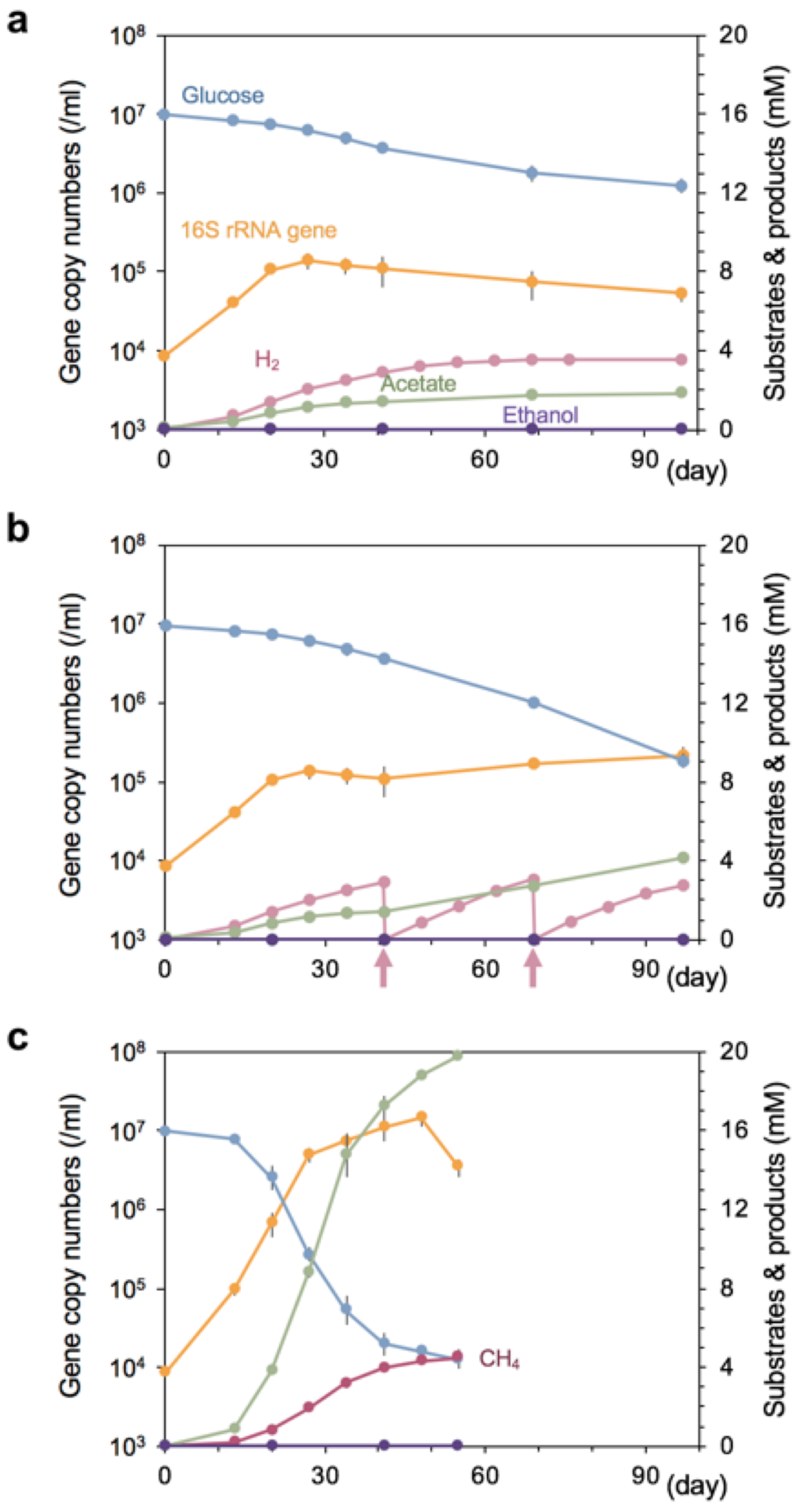


**Fig. S7.** Growth and metabolic behavior of RT761 showing the effects of $H_2$ removal on growth. (**a**) Pure culture. (**b**) Pure culture with periodic purging of culture vial head space with $N_2/CO_2$ (arrow). (**c**) Co-culture with $H_2$-scavenging methanogenic archaeon, *Methanothermobacter thermoautotrophicus* str. ΔH. Means and standard deviation (error bars) of triplicate cultures are shown. In all culture conditions, ethanol was detected in HPLC analysis, but its concentration was too low (at least lower than 0.2 mM) for determining accurate concentrations.

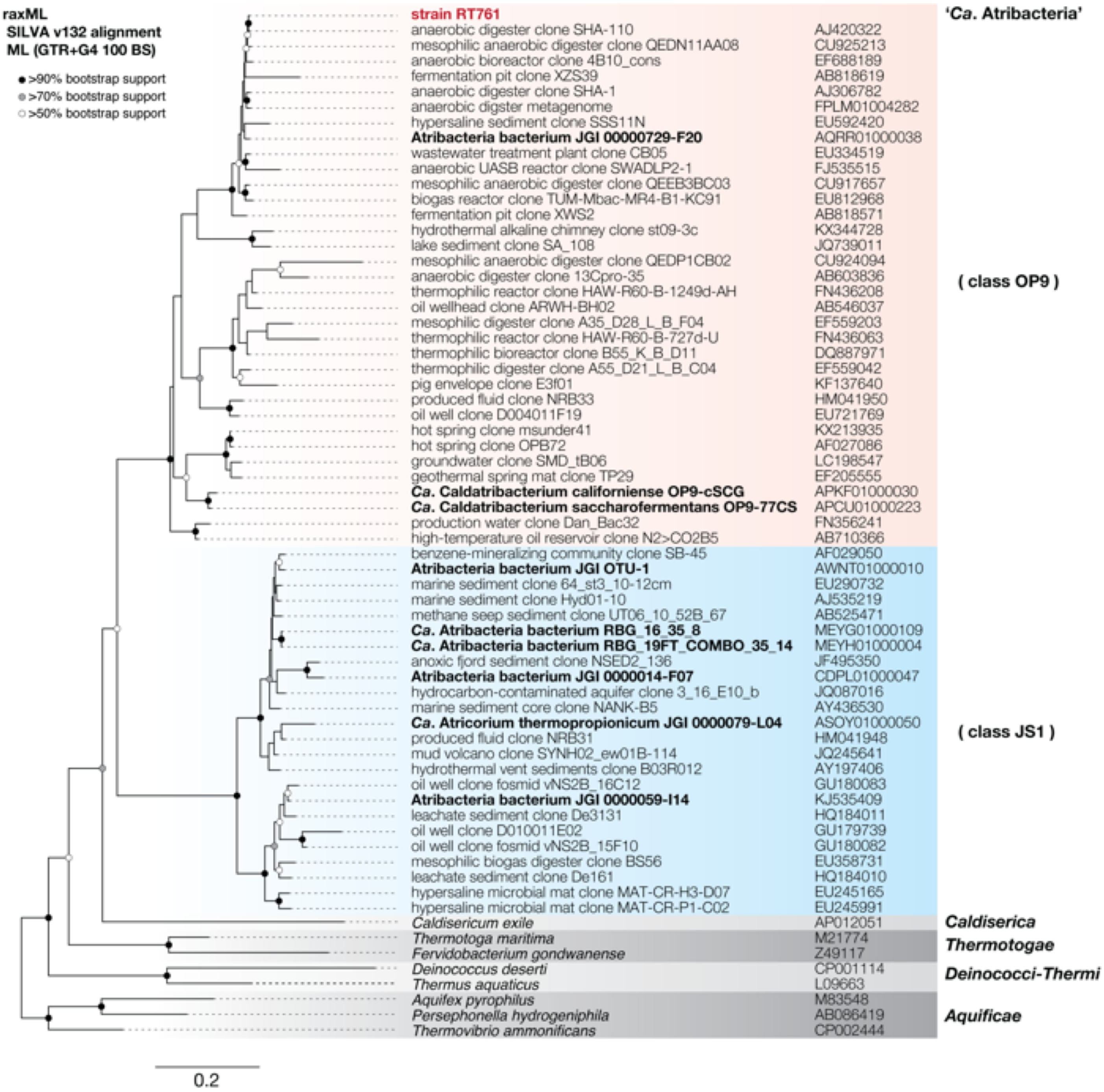


**Fig. S8.** A 16S rRNA gene phylogenetic tree (Maximum likelihood tree) showing the relationship between strain RT761 (red) and relatives including single-cell genomes and metagenomic bins (bold) assigned to '*Ca*. Atribacteria' classes OP9 (red background) and JS1 (blue background). Bootstrap values greater than 50% (white circle), 70% (gray), and 90% (black) are shown.

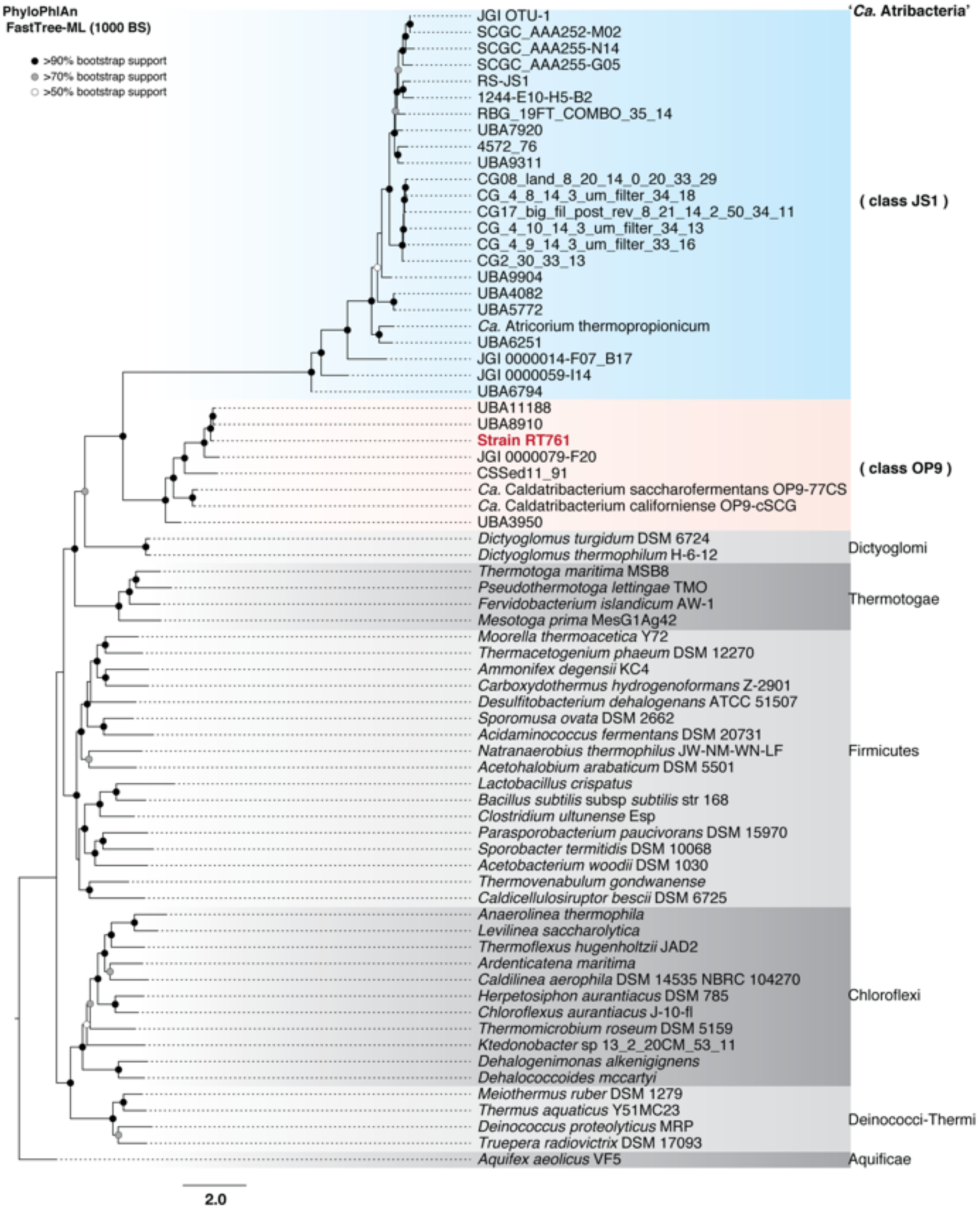


**Fig. S9.** Phylogenomic analysis of strain RT761 (red) and relatives using a concatenated alignment of conserved marker genes (PhyloPhlAn). Genomes were selected for '*Ca.* Atribacteria' classes classes OP9 (red background) and JS1 (blue background) and other related phyla. Bootstrap values greater than 50% (white circle), 70% (gray), and 90% (black) are shown.

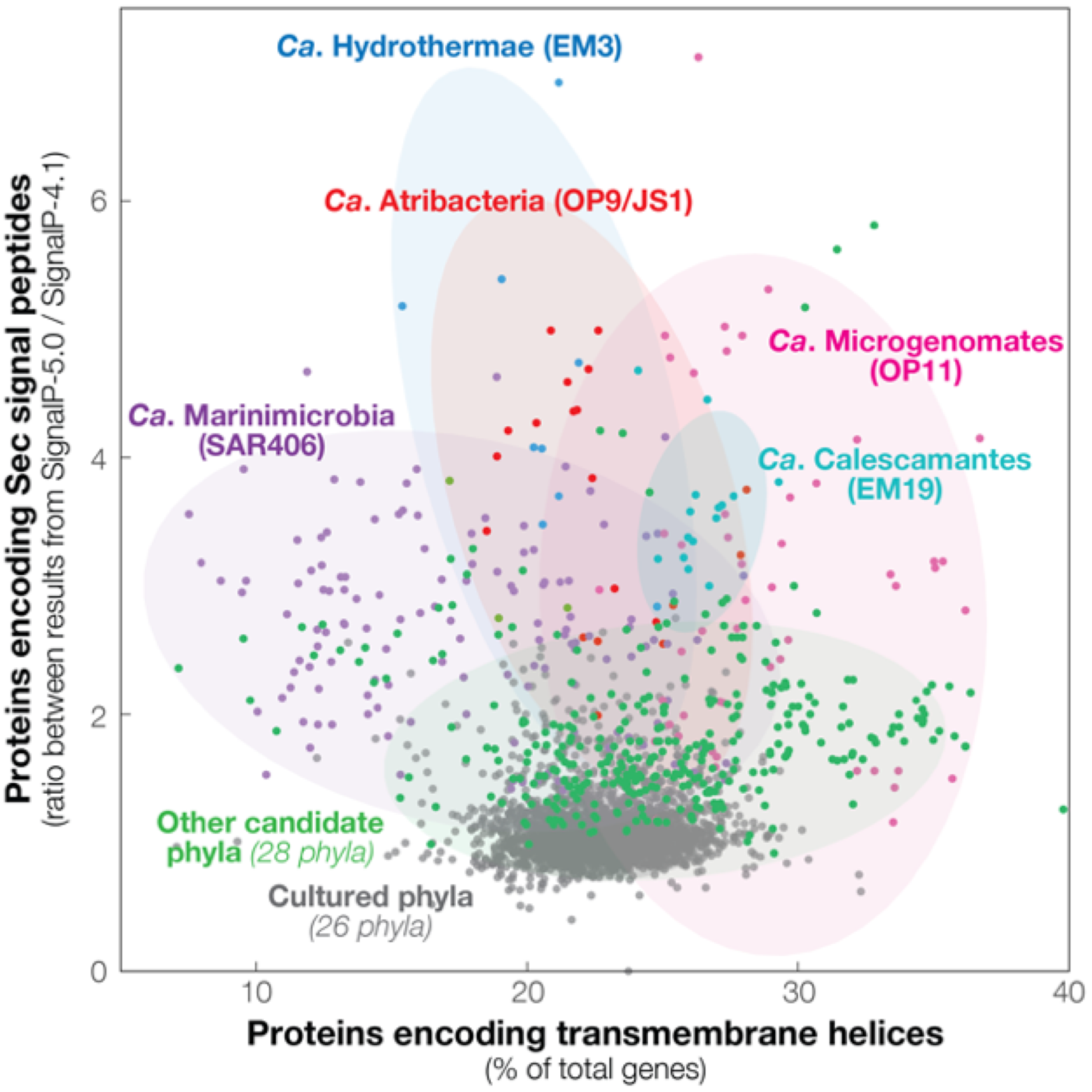


**Fig. S10.** Genomic compositions of membrane-related features in yet-to-be-cultured candidate phyla. The horizontal axis shows the genomic proportion proteins encoding transmembrane helices. The vertical axis shows the ratio of proportions of proteins encoding Sec signal peptides estimated by SignalP-5.0 and SignalP-4.1. Genomes of '*Ca.* Atribacteria' (OP9 and JS1; red), '*Ca.* Microgenomates' (OP11; pink), '*Ca.* Calescamantes' (EM19; light blue), '*Ca.* Hydrothermae' (EM3; blue), '*Ca.* Marinimicrobia' (SAR406; purple), 28 other candidate phyla (green), and 26 cultured phyla (gray, see Fig. S6) are plotted. Confidence ellipses (95%) are shown for each aforementioned group.

**Supplementary Discussions**

**1. Localization of ribosomes in RT761 cells**

In this experiment, whether the location of ribosome is corresponding to the region inside the intracytoplasmic membrane (ICM) is not clear because membrane lipid staining could not apply to formamide-fixed cells, but is supported by microscopic observations: Fluorescence signal of stained rRNA was almost absent in cell poles, which entirely coincided with the cytoplasmic membrane-bounded space (CBS) (Fig. 2). Additionally, transmission electron micrographs show that the ribosome-like, electron-dense particles were mainly observed in ICM-bound space (IBS) (Fig. 1 and Supplementary Fig. S1).

**2. Unique Sec-secreted proteins in the RT761 genome**

SignalP-5.0 estimated 2.67 times more Sec-secreted proteins than SignalP-4.0 estimated in RT761 genomes. Evaluation of all gram-negative type strain genomes revealed that 26 out of 29 phyla have consistent predictions between SignalP-4.1 and SignalP-5.0 (1.1 ± 0.2 [S.D.] times more in SignalP-5.0 on average). Note that the reference databases used by SignalP-4.1 and SignalP-5.0 are similar in phylogenetic composition and diversity (Supplementary Table S4). Through further analysis of genomes from 33 candidate phyla, we discover that only two candidate phyla, '*Ca.* Calescamantes' (EM3) and '*Ca.* Microgenomates' (OP11) within '*Ca.* Patescibacteria', have RT761-like signatures (Supplementary Fig. S10) (1.8± 0.6 [S.D.] times more in SignalP-5.0 on average for other candidate phyla), indicating that the above features of RT761 are not an artifact from lack of *Ca.* Atribacteria-derived sequences in the SignalP reference databases. Comparison of hydrophobicity, signal peptide length, and number of NT cationic residues interestingly revealed no differences between Sec-secreted proteins predicted by SignalP-5.0 and SignalP-4.1, suggesting that the Sec-secreted proteins of RT761 may have unique signal peptides that could only be predicted through integration of a recurrent neural network implemented in SignalP-5.0.
